# Receptor binding and proteolysis do not induce large conformational changes in the SARS-CoV spike

**DOI:** 10.1101/292672

**Authors:** Robert N. Kirchdoerfer, Nianshuang Wang, Jesper Pallesen, Daniel Wrapp, Hannah L. Turner, Christopher A. Cottrell, Kizzmekia S. Corbett, Barney S. Graham, Jason S. McLellan, Andrew B. Ward

**Author notes:** Corresponding author ABW. Current address: Department of Molecular Biosciences, The University of Texas at Austin, Austin, TX, 78712.

## Abstract

Severe acute respiratory syndrome coronavirus (SARS-CoV) emerged in 2002 as a highly transmissible pathogenic human betacoronavirus. The viral spike glycoprotein (S) utilizes angiotensin-converting enzyme 2 (ACE2) as a host protein receptor and mediates fusion of the viral and host membranes, making S essential to viral entry into host cells and host species tropism. As SARS-CoV enters host cells, the viral S undergoes two proteolytic cleavages at S1/S2 and S2’ sites necessary for efficient membrane fusion. Here, we present a cryo-EM analysis of the trimeric SARS-CoV S interactions with ACE2 and of the trypsin-cleaved S. Surprisingly, neither binding to ACE2 nor cleavage by trypsin at the S1/S2 cleavage site impart large conformational changes within S or expose the secondary cleavage site, S2’. These observations suggest that S2’ cleavage does not occur in the S prefusion conformation and that additional triggers may be required.

---

Severe acute respiratory syndrome coronavirus (SARS-CoV) emerged in humans in 2002 and rapidly spread globally causing 8,096 cases and 774 associated deaths in 26 countries through July 2003^1^. SARS-CoV reappeared in a second smaller outbreak in 2004, but has since disappeared from human circulation. However, closely related coronaviruses, such as WIV1, currently circulate in bat reservoirs and are capable of utilizing human receptors to enter cells^2^. The more recent emergence of Middle East respiratory syndrome coronavirus (MERS-CoV)^1^ and the likelihood of future zoonotic transmission of novel coronaviruses to humans from animal reservoirs make understanding the coronavirus infection cycle of great importance to human health.

Coronaviruses are enveloped viruses possessing large, trimeric spike glycoproteins (S) required for the recognition of host receptors for many coronaviruses as well as the fusion of viral and host cell membranes for viral entry into cells^3^. During viral egress from infected host cells, some coronavirus S proteins are cleaved into S1 and S2 subunits. The S1 subunit is responsible for host-receptor binding while the S2 subunit contains the membrane-fusion machinery. During viral entry, the S1 subunit binds host receptors in an interaction thought to expose a secondary cleavage site within S2 (S2’) adjacent to the fusion peptide for cleavage by host proteases^4–7^. This S2’ proteolysis has been hypothesized to facilitate insertion of the fusion peptide into host membranes after the first heptad repeat region (HR1) of the S2 subunit rearranges into an extended α-helix^8–10^. Subsequent conformational changes in the second heptad repeat region (HR2) of S2 form a six-helix bundle with HR1, fusing the viral and host membranes and allowing for release of the viral genome into host cells. Coronavirus S is also the target of neutralizing antibodies^11^, making an understanding of S structure and conformational transitions pertinent for investigating S antigenic surfaces and designing vaccines.

The SARS-CoV S1 subunit is composed of two distinct domains: an N-terminal domain (S1 NTD) and a receptor-binding domain (S1 RBD) also referred to as the S1 CTD or domain B. Each of these domains have been implicated in binding to host receptors, depending on the coronavirus in question. However, most coronaviruses are not known to utilize both the S1 NTD and S1 RBD for viral entry^12^. SARS-CoV makes use of its S1 RBD to bind to the human angiotensin-converting enzyme 2 (ACE2) as its host receptor^13,14^.

Recent examination using cryo-electron microscopy (cryo-EM) has illuminated the prefusion structures of coronavirus spikes^15–22^. Initial examination of HCoV-HKU1 S showed that the receptor-binding site on the S1 RBD was occluded when the RBD was in a ‘down’ conformation and it was hypothesized that conformational changes were required to access this site^16^. Subsequent studies of the highly pathogenic human coronavirus S proteins of SARS-CoV^15,22^ and MERS-CoV^17,22^ showed that these viral S1 RBD do indeed sample an ‘up’ conformation where the receptor-binding site is accessible. These structural studies also located the positions of the S1/S2 and S2’ cleavage sites on the prefusion spike. The S1/S2 site lies within a surface exposed loop in the second subdomain of S1^16^. However, the S2’ site lies closer to base of the spike and though this region is located on the surface of the spike, cleavage at this site is prevented by surrounding protein elements^17^.

To examine the hypothesized conformational transitions induced by proteolysis and receptor binding, we used single-particle cryo-EM to determine structures of S in uncleaved, S1/S2 cleaved and ACE2-bound states. Three-dimensional classification of the S1 RBD positions and corresponding atomic protein models revealed that neither ACE2-binding nor trypsin cleavage at the S1/S2 boundary induced substantial conformational changes in the prefusion protein conformation to expose the S2’ proteolysis site, suggesting additional requirements for fusion-associated conformational changes and secondary protease cleavage.

## Results

### Structural description of SARS-CoV S 2P ectodomain

To take advantage of improved protein expression and increased prefusion conformational stability, the SARS-CoV S ectodomain used for cryo-EM studies contained two proline substitutions (2P) in the S2 region^17^. These mutations had similar prefusion stabilizing effects on MERS-CoV S where their inclusion did not alter presentation of known epitopes or receptor recognition compared with wild-type MERS-CoV S. To establish a baseline for any potential conformational changes, we determined structures of the SARS-CoV S 2P ectodomain. This included a 3.2 Å resolution C3-symmetric reconstruction and several asymmetric reconstructions (3.9-4.5 Å, see below) (Fig. 1 and Supplementary Table 1 and 4 and Supplementary Fig. 1). The S1 NTD and some of the S1 RBD, while clearly present, appear poorly resolved in the cryo-EM maps, precluding a *de novo* build of these regions. For this reason, available SARS-CoV S crystal structures^22,23^ of corresponding domains were placed into these densities. Overall, the SARS-CoV S 2P structure resembles that of other coronaviruses, particularly those of the betacoronavirus genus^16,17,19,22^ and the previously published SARS-CoV wild-type prefusion ectodomains^15,22^. The S1 domains surround the helical S2 subunits and interdigitate at the membrane-distal apex of the trimeric spike. The structure of the S2 subunit appears to be highly conserved across coronavirus genera^18,20,21^. The SARS-CoV prefusion wild-type and 2P structures are nearly identical, including regions proximal to the 2P mutation site (Fig. 1b and c).

**Figure 1:**
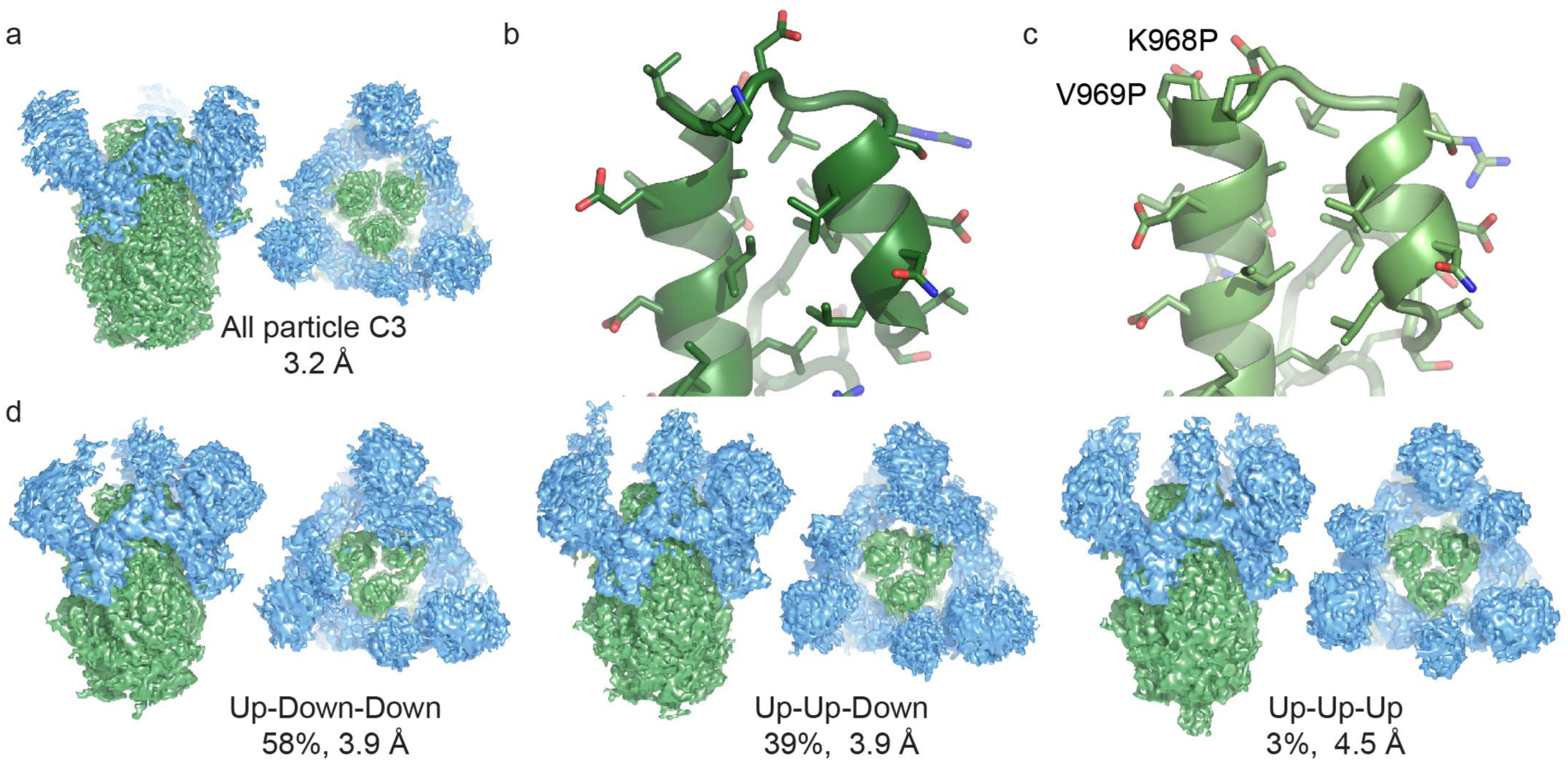
Structure of the SARS-CoV S 2P ectotodomain. a) The C3 symmetrized reconstruction of all particles within the dataset resembles that of other betacoronaviruses. Side and membrane-distal top views (90˚ rotation about x-axis) are shown. Coordinate models derived from cryo-EM reconstructions of b) the wild-type SARS-CoV S ectodomain^22^ and c) prefusion stabilized SARS-CoV S 2P ectodomain adopt identical conformations near the 2P mutation site. Coordinate models derived from C3 symmetry cryo-EM reconstructions are shown. d) Classification of heterogeneity within the S1 RBD reveals a distribution of RBD configurations with the single-‘up’ conformation being most prevalent. S1 regions are shown in blue and S2 regions are shown in green.

Although the overall structures of SARS-CoV S are similar, we were able to build several S2 protein regions excluded from the previous SARS-CoV models, including a fusion-peptide-adjacent region as well as additional amino acids in the S2 C-terminal connector domain. The region C-terminal to the fusion peptide, known as FP2, has been suggested to act as the second part of a bipartite fusion peptide for SARS-CoV S^24^. Comparison of this FP2 region in the structures presented here with the MHV^19^, HCoV-HKU1^16^, MERS-CoV^17,22^, HCoV-NL63^20^ and Porcine deltacoronavirus^18,21^ S structures indicates that SARS-CoV adopts a unique conformation in this region (Fig. 2). Whereas most other coronaviruses possess two short α-helices with a cross-linking disulfide bond, in SARS-CoV a similar disulfide bond bridges the two strands of a beta-hairpin. The unique SARS-CoV S conformation of FP2 and the lack of cross-group sequence conservation beyond the disulfide bond in this region suggest that SARS-CoV may use a distinct mechanism of FP2 membrane insertion.

**Figure 2:**
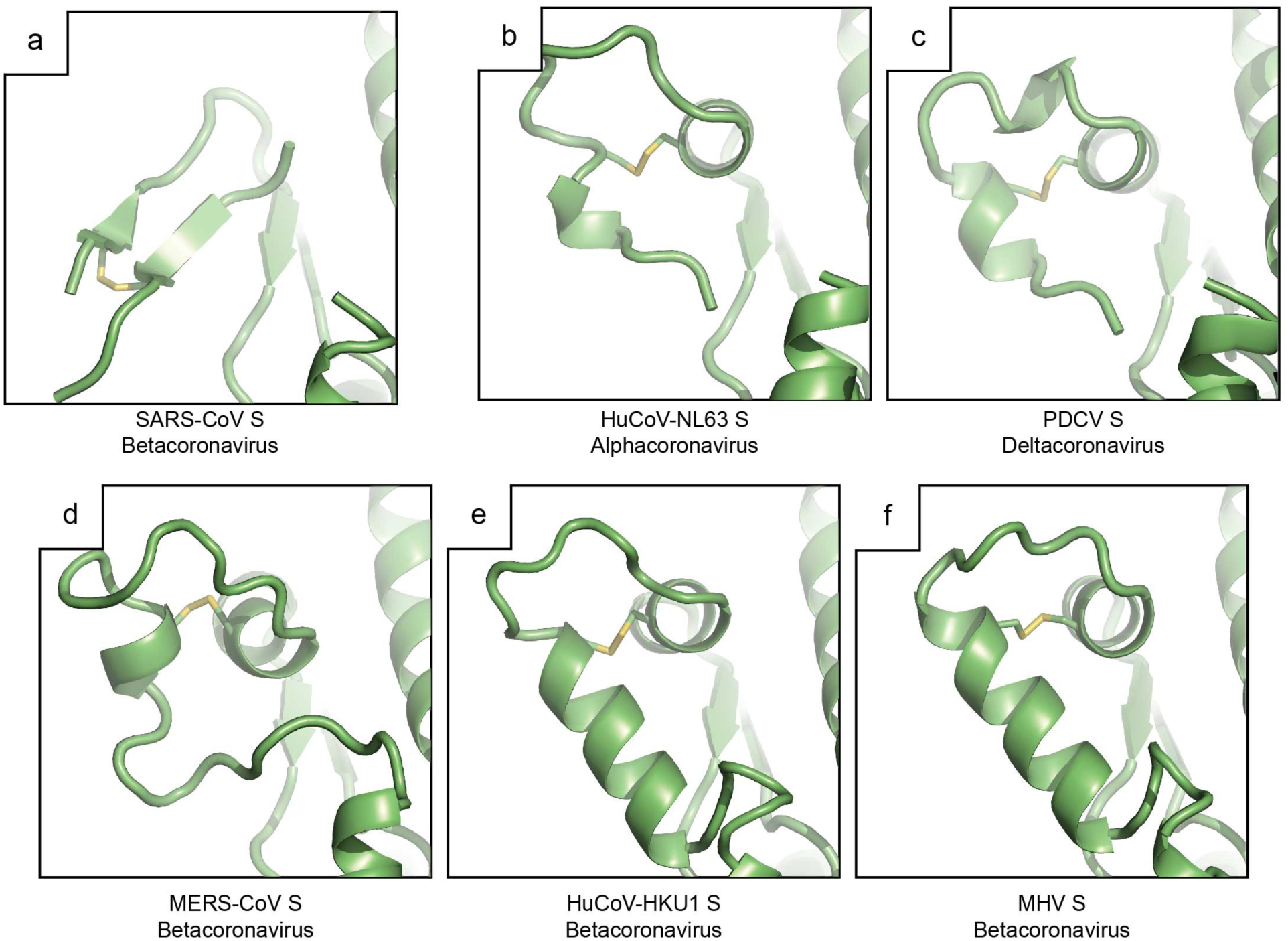
Comparison of putative bi-partite fusion peptide regions. a) The SARS-CoV S FP2 region of the bi-partite fusion peptide^24^ adopts a conformation distinct from equivalent regions in b) alpha-(HuCoV-NL63, 5SZS.pdb^20^) and c) deltacoronavirus spikes (PDCV S, 6BFU.pdb^21^) as well as other d-f) betacoronaviruses (MERS-CoV S, 5W9I.pdb^17^, HuCoV-HKU1 S, 5I08.pdb^16^, MHV S, 3JCL.pdb^19^).

**Figure 3:**
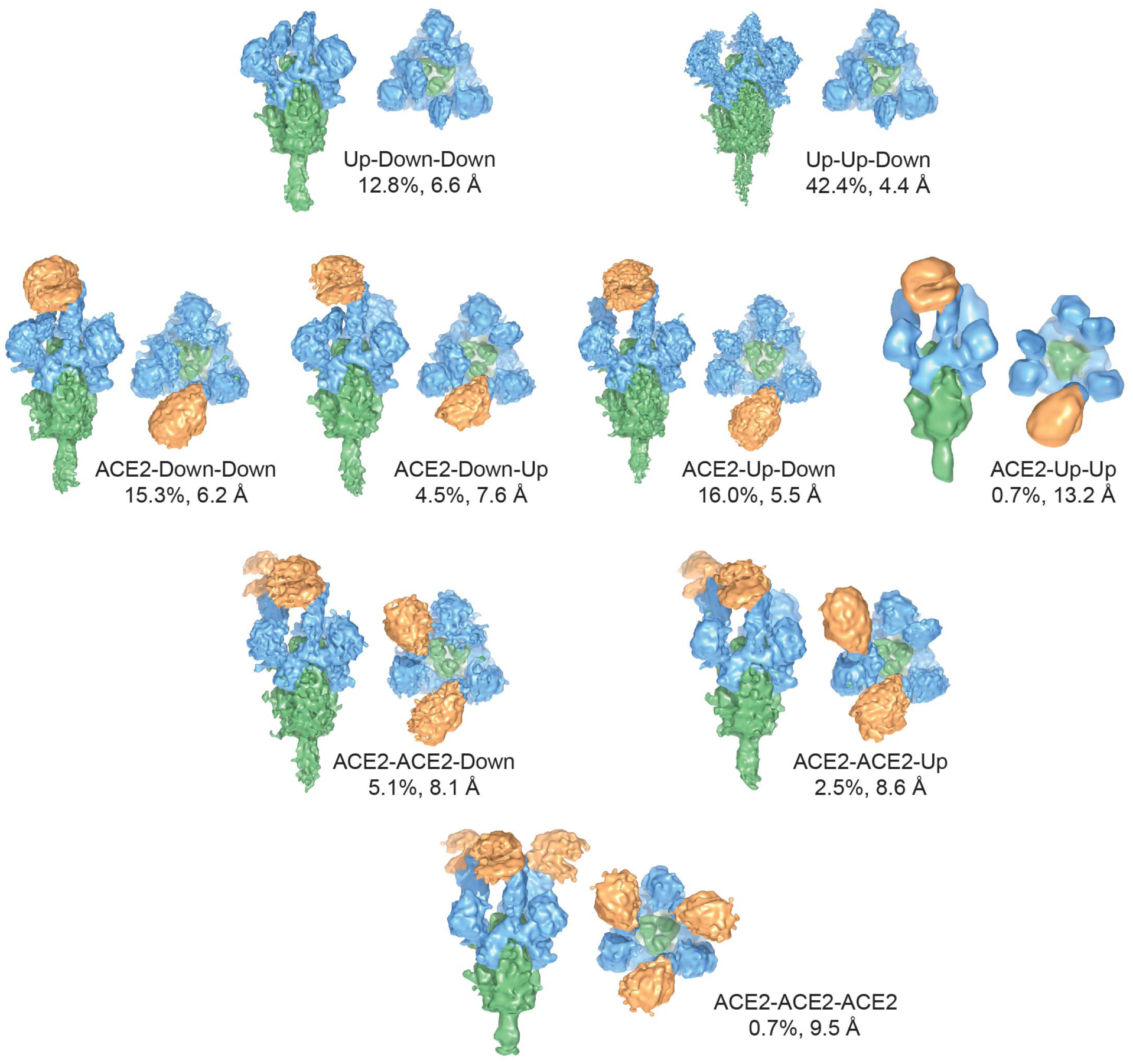
Compositional and conformational heterogeneity of ACE2-bound SARS-CoV spikes. Classification of heterogeneity within the SARS-CoV S – ACE2 complex reveals that 55% of the particles are not bound by ACE2. Of those particles which are bound by ACE2, each configuration of the S1 RBD is observed with classes containing triple ‘up’ conformation S1 RBD being the most poorly represented. The labeled description of each class begins at the lower RBD when viewed at the membrane distal apex and proceeds clockwise. Side views and membrane-distal top views are shown for each reconstruction. S1 regions are shown in blue, S2 regions are shown in green and soluble ACE2 is shown in orange.

As observed in the previous SARS-CoV and MERS-CoV S structures^15,22^, the trimeric S adopts two distinct conformations related to each of the S1 RBD. The ‘down’ conformation caps the S2 helices and makes extensive contacts with the S1 NTD. The ‘up’ conformation of the S1 RBD exposes the S1 RBD receptor-binding site. It has been previously reported that for wild-type SARS-CoV S, 56% of the particles contained three ‘down’ RBD conformations while 44% contained a single ‘up’ S1 RBD conformation^22^. To examine the conformation of the S1 RBD among our SARS-CoV S 2P ectodomains, we used a local masking and 3-D sorting strategy^17^ to more accurately classify the conformations as being either ‘down’ or ‘up’ at each of the three S1 RBD positions within the trimer. This analysis revealed that the majority of the S 2P proteins were in the single-‘up’ conformation (58%) with lesser amounts of double- and triple-‘up’ conformations (39% and 3% respectively) and with no all-‘down’ conformation observed. The increased propensity to adopt the ‘up’ S1 RBD conformation may indicate a difference in the coronavirus S containing the 2P mutations, however other differences in sample preparation cannot be ruled out.

### ACE2 and S1 C-terminal domains

To examine the structure of SARS-CoV S bound to its receptor, ACE2, we combined SARS-CoV S 2P ectodomain with an excess of soluble human ACE2 with subsequent purification by size-exclusion chromatography and immediate cryo-EM specimen preparation. Initial sorting of particle heterogeneity indicated spikes could be split into ACE2-bound (45%) and unbound (55%) classes. Using a similar masking and 3-D sorting strategy as above we sorted the unbound S class further into classes with S1 conformations of one or two ‘up’ S1 RBDs (Fig 3 and Supplementary Tables 2-4 and Supplementary Fig. 2-4). We did not observe an all-‘down’ class nor a three ‘up’ S1 RBD class indicating a low prevalence of these conformations among the unbound spikes. Expanding our 3D sorting strategy, we classified our ACE2-bound particles at each S1 RBD position and identified single, double and triple ACE2-bound S. We were further able to identify S1 RBD conformations at the non-ACE2 occupied RBD positions to represent each population of S1 RBD conformations among ACE2-bound S.

As hypothesized by previous structural work^15–17,22^, the S1 RBD recognizes ACE2 with an ‘up’ S1 RBD conformation. The proportion of total ‘up’ S1 RBD conformations within the ACE2-bound and -unbound classes is nearly identical within this dataset (58% ‘up’ S1 RBD), similar to the proportion of total ‘up’ S1 RBD in the SARS S 2P ectodomain dataset (48%). This strongly suggests that binding of a single ACE2 receptor does not induce adjacent S1 RBDs to transition from a ‘down’ to ‘up’ conformation. Hence, ACE2 is more likely to bind to an already ‘up’ S1 RBD rather than inducing the conformational changes that are required for the S1 RBD to become accessible to ACE2.

It is noteworthy that despite prolonged co-incubation and an excess of ACE2, we had difficulties in saturating the S1 RBD with ACE2 in the context of trimeric S ectodomain. This poor saturation is illustrated by the small proportion of triple-bound ACE2 and the majority of spikes that are unbound by receptor. In contrast, isolated recombinant S1 RBD easily binds ACE2 and is capable saturating ACE2 on target cells to block S-mediated entry^14^. Our observed sub-stoichiometric ACE2 binding to trimeric spikes is consistent with the difficulty in using soluble ACE2 receptor to neutralize SARS-CoV S pseudotyped onto VSV^25^. The reduced binding of ACE2 to trimeric spikes is likely due to the incomplete exposure and conformational flexibility of the S1 RBD. Incomplete neutralization with soluble receptor was not encountered for MHV which binds CEACAM1a via its S1 NTD, which does not undergo conformational changes^19,26^.

Similar to recently published MERS-CoV S structures^17^, the ACE2-bound RBD adopts a much more extended and rotated conformation compared to S1 RBD modeled in previous SARS-CoV S structures^22^. This difference is likely due to poor density in the hinge regions between the S1 RBD and subdomain 1 (SD-1) in these previous reconstructions^15,22^ rather than the presentation of a unique receptor-bound conformation. Indeed, the bound ACE2 receptor and S1 RBD for all reconstructions here show poorer density quality than the less mobile regions of the SARS-CoV S (Fig 4). To improve the density for ACE2-bound S1 RBD, we used focused refinement on this region to overcome the flexibility of these domains relative to the rest of S. This yielded a 7.9 Å resolution reconstruction with improved local density quality (Fig 4b and c). We successfully placed the crystal structure of the SARS-CoV S1 RBD bound to ACE2 (2AJF.pdb^23^) into this density as a rigid body indicating that the previously determined crystal structure accurately recapitulates the conformation between the ACE2-bound S1 RBD in the trimeric spike.

**Figure 4:**
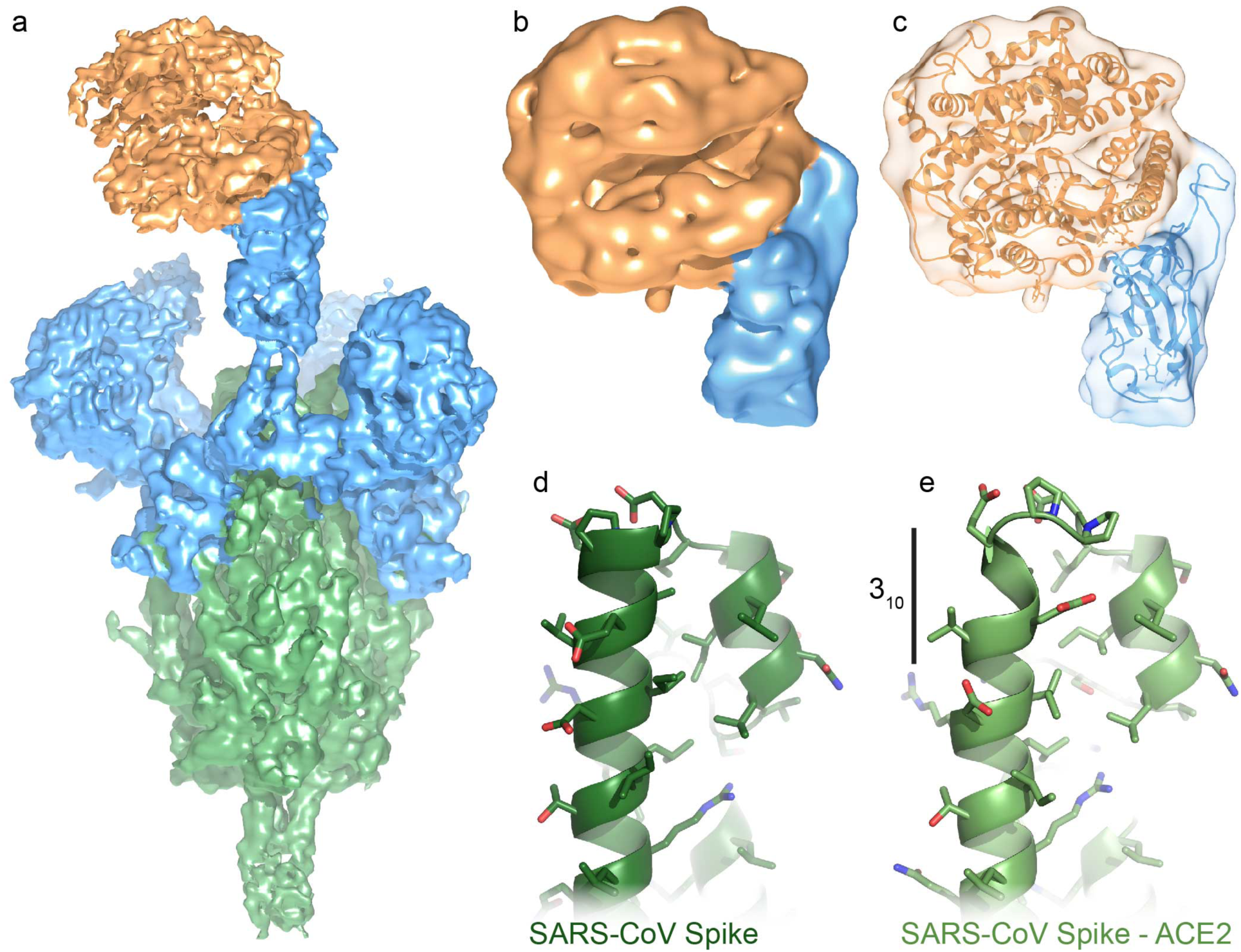
ACE2-receptor binding and induced conformational changes. a) ACE2 binds the SARS-CoV S1 RBD in an ‘up’ conformation. b) Focused refinement was used on the S1 RBD – ACE2 portion of the particles to improve the density of this flexible region. c) Rigid-body fitting of the SARS-CoV S1 RBD – ACE2 complex crystal structure (2AJF.pdb^23^) indicates that the crystal structure is representative of the ACE2-bound trimeric spike. Comparison of the d) SARS-CoV S 2P central helices with the e) SARS-CoV S 2P central helix which has been uncapped by an ACE2-bound S1 RBD demonstrates a transition to a short 3_10_ helix. Colors are as in Fig. 3.

The ACE2-bound, S1 RBD extends upwards and rotates away from contacts with nearby amino acids. Hence, any conformational changes induced by receptor binding to the S1 RBD are more likely to be caused by the absence of the S1 RBD contacts in the ‘up’ conformation, rather than the formation of additional contacts (Supplemental Figure 5). This model provides a flexible mechanism for how different coronavirus spikes can bind to different protein receptors with their S1 RBD and facilitate fusion with host cells. Moreover, movements of the S1 RBD to the ‘up’ conformation disrupt molecular interactions with the S2 central α-helices which may play a key role in receptor-induced conformational changes leading to membrane fusion.

### SARS-CoV S2 fusion machinery and receptor binding

Several biochemical and virological studies have suggested that binding of a host protein receptor to coronavirus spikes induces conformational changes leading to fusion and exposure of the S2’ protease site for cleavage^4,6^. Comparison of our ACE2-bound 2P-stabilized spike structures with those of both the wild-type SARS-CoV S and the 2P-stabilized SARS-CoV S indicate that overall, these structures are surprisingly similar and do not exhibit large conformational differences. The absence of conformational changes includes regions near the S2’ cleavage site, fusion peptide and S2 heptad repeat 1 (HR1). However, fine examination of the ACE2-bound S structure revealed a modest conformational change in the S2 central helix where the S2 had been uncapped by a receptor-bound RBD. In both the wild-type^15,22^ and 2P unbound structures, the central helix presents as an α-helix. However, in the ACE2-bound structure, the upper portion of this helix has transitioned to a 3_10_-helix (Fig. 4d and e). This subtle transition may indicate that conformational changes towards fusion are initiated in the central helix. N-terminal to the central helix are the 2P mutations that stabilize the spike in its prefusion conformation. These stabilizing mutations may block the propagation of conformational changes beyond the subtle change to a short 3_10_-helix, providing a possible mechanism for their prefusion stabilizing effect.

### Trypsin-cleaved SARS-CoV spike

Unlike some other coronavirus spikes such as MERS-CoV S, SARS-CoV S lacks an amino acid sequence capable of recognition by host furin proteases at the S1/S2 cleavage site. However, SARS-CoV S can be cleaved *in vitro* by exogenous trypsin at an equivalent site, and it has been proposed that a similar cleavage event may occur *in vivo* by trypsin-like proteases during viral entry^4,27^. Moreover, trypsin cleavage of the SARS-CoV S potentiates S for more efficient membrane fusion^27,28^. For MERS-CoV S, cleavage at S1/S2 enables more efficient receptor recognition^6^. In addition, coronaviruses possess a secondary cleavage site in S2 called S2’^27^. This site is thought to be cleaved by host proteases after receptor recognition^4–6^ and liberates the viral fusion peptide to insert into host membranes in a S2 pre-hairpin intermediate^29^. The S2’ cleavage has been proposed to occur after an arginine residue conserved across coronavirus genera (SARS-CoV S Arg797)^30^.

To examine the trypsin protease sensitivity of our coronavirus spikes, we carried out limited proteolysis experiments of both wild-type and 2P SARS-CoV S ectodomains in the presence and absence of soluble ACE2 receptor. A time course of the proteolysis revealed that all four samples are cleaved to S1 and S2 products at equivalent rates at similar sites (Supplementary Fig. 6). Nearing the end of the time course additional lower molecular weight bands are observed which we interpret to be degradation of the S1 subunit. Regardless of which construct was used or whether ACE2 was bound to the S ectodomains, there is no prominent band that corresponds to a S2’ cleavage product (approximately 52 kDa).

To analyze the cleavage products in detail, we performed cryo-EM analysis on the trypsin-cleaved SARS-CoV S 2P ectodomain. Using all-particles and C3 symmetry yielded a reconstruction at 3.3 Å resolution (Fig. 5, Supplementary Tables 1 and 4 and Supplementary Fig. 7). The short loop containing the S1/S2 cleavage site is disordered in the uncleaved spike reconstruction and remains disordered in the trypsin cleaved reconstruction. Moreover, examination of the structure models indicates no significant differences between the trypsin-cleaved and uncleaved SARS-CoV S (Fig. 5b). Fine sorting of S1 RBD positions of the trypsin-cleaved S reveals a very similar distribution of ‘up’ S1 RBD conformations available for receptor binding as in the uncleaved samples, although we additionally observe a small proportion of S1 RBD in the all-‘down’ conformation (Fig. 5c). These results indicate that trypsin-cleavage at S1/S2 does not impart large conformational changes on the SARS-CoV S and justifies the removal of S1/S2 cleavage sites for the production of more homogeneous material as vaccine immunogens. This suggests that although cleavage at S1/S2 may remove an obstacle for conformational changes leading to fusion, S1/S2 cleavage alone does not produce significant conformational changes.

**Figure 5:**
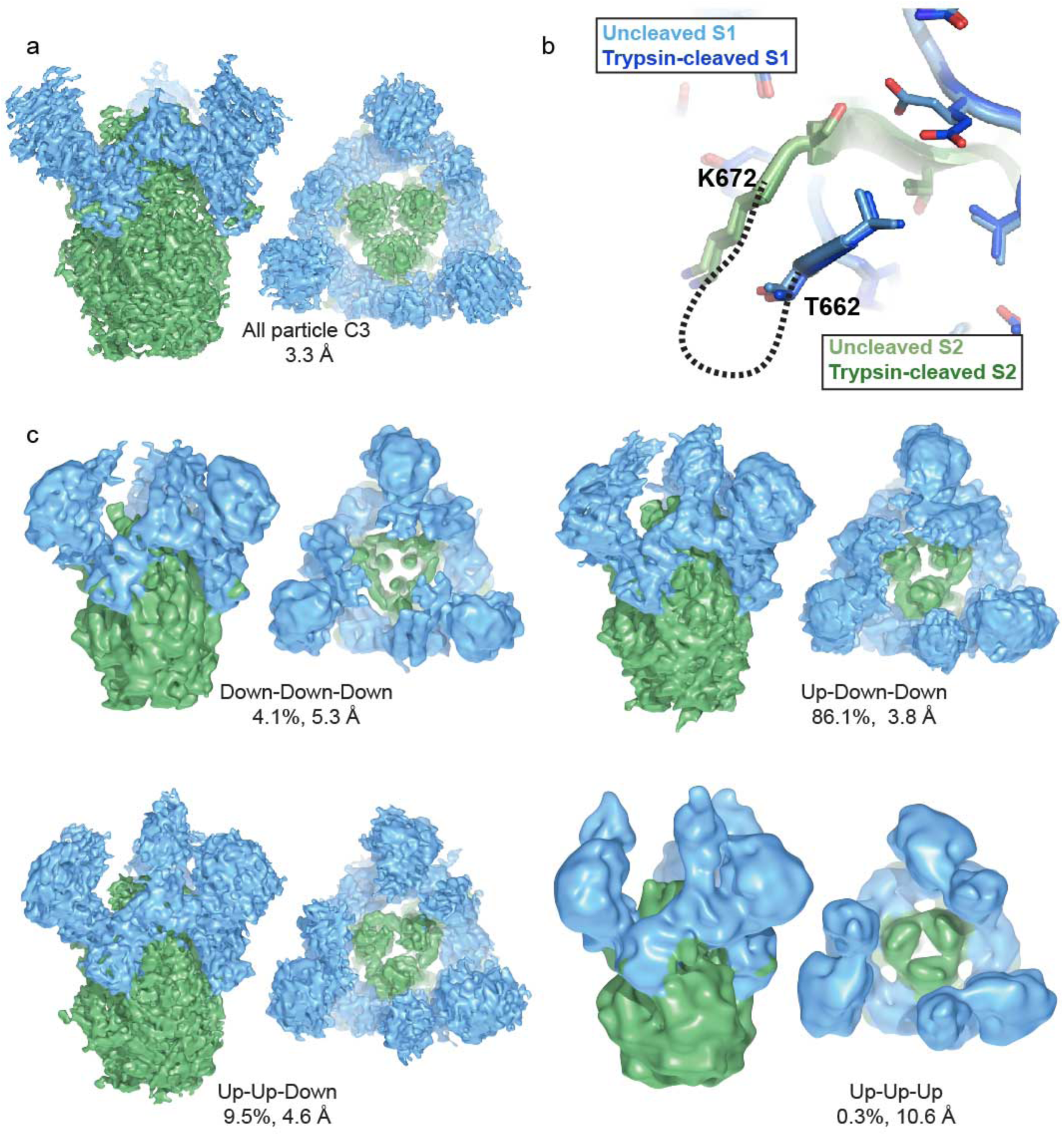
Trypsin cleavage does not impart conformational changes in prefusion SARS-CoV spike. a) An all particle reconstruction of trypsin-cleaved SARS-CoV S 2P with C3 symmetry. b) Comparison of the trypsin-cleaved and uncleaved SARS-CoV S 2P coordinate models reveals no differences in protein structure near the cleavage site. The last residue in S1 (T662) and the first residue in S2 (K672) visualized in the coordinates are labeled. c) Classification of the heterogeneous S1 RBD in the trypsin-cleaved S shows a predominance of the single-‘up’ conformation as well as the observation of an all-‘down’ conformation not seen in the uncleaved S 2P sample. Colors are as in Fig. 1.

In agreement with the *in vitro* limited proteolysis experiments, the S2’ site remains uncleaved in the trypsin-cleaved spike ectodomain structures. The arginine at the proposed S2’ cleavage site in all coronavirus spike structures published to date and presented here is blocked from cleavage by a loop protruding from just N-terminal to the fusion peptide and by an N-terminal helix of S2 HR1 (Fig 6). Exposure of this site for cleavage may require remodeling of this penultimate loop or HR1 beyond the conformation observed in the prefusion state. We hypothesize that additional triggers beyond cleavage at the S1/S2 site or protein-receptor binding are needed to transition the spike from its prefusion state to a yet to be observed intermediate. Further, this intermediate may be the target for S2’ cleavage before final adoption of the postfusion six-helix bundle conformation and membrane fusion. Indeed, it has been suggested that for some strains of MHV, the spike undergoes a S2’-like cleavage after transitioning to a SDS-resistant trimer in the presence of its protein receptor^4^.

**Figure 6:**
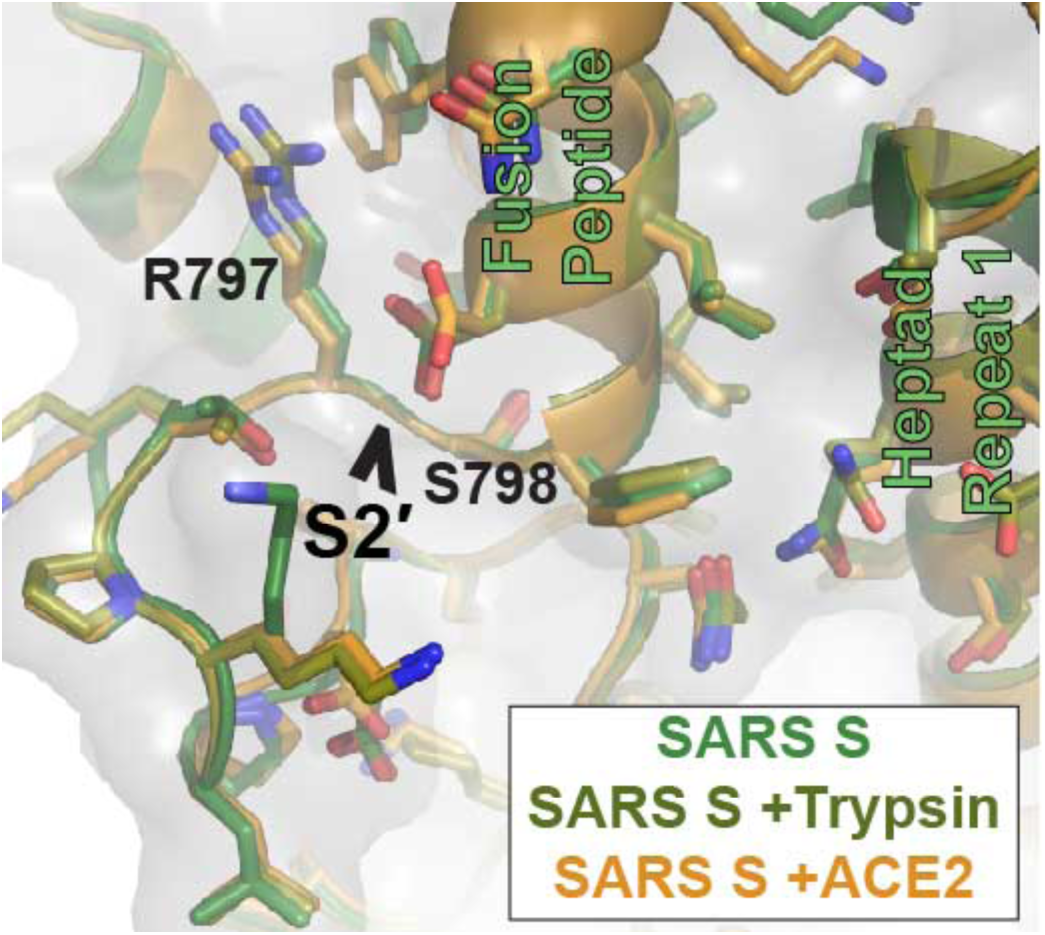
S2’ secondary cleavage site is occluded. Superimposition of the SARS-CoV S 2P, uncleaved, cleaved and ACE2-bound structures reveals no structural changes within this region. The S2’ site after amino acid R797 is shielded from cleavage by an adjacent loop and the fusion peptide which packs against an N-terminal region of S2 HR1.

## Discussion

SARS-CoV is a highly transmissible pathogen, which emerged from an animal reservoir and rapidly spread around the world^1^. Though eventually removed from human circulation through public health measures, SARS-like coronaviruses continue to circulate in animal reservoirs^2,31^ and there are no vaccines or virus-specific treatments available for human use. Understanding how coronavirus S glycoproteins are processed and bind to host receptors is key to the development of coronavirus vaccines and therapeutics.

Here, we present the first structures of a trimeric coronavirus spike ectodomain bound to a protein receptor as well as the first structures of prefusion S1/S2 cleaved coronavirus spikes. Our structures of the SARS-CoV S 2P ectodomain bound to a soluble form of human ACE2 receptor show that any conformational changes induced in S by receptor binding are more likely to be due to the disruption of protein-protein interactions rather than the formation of additional contacts between the S1 RBD and other regions of S (Supplemental Fig. 5). The only conformational change that we observe in the ACE2-bound structures is the transition to a short 3_10_-helix at the top of the S2 central α-helix when uncapped by receptor-bound S1 RBD suggesting that more extensive conformational changes may be initiated here. The 2P prefusion stabilizing mutations adjacent to this region may block more extensive transitions from occurring. However, both wild-type and 2P SARS-CoV S ectodomains bound to ACE2 were cleaved equally by trypsin, indicating that at least at the level of protease susceptibility, these two receptor-bound complexes are equivalent.

Our examinations of both receptor-bound and trypsin-cleaved prefusion SARS-CoV S proteins indicate that neither receptor binding nor trypsin cleavage induces large conformational changes in S and do not expose the S2’ cleavage site for proteolysis. These data suggest that SARS coronavirus spikes may require an additional factor to undergo such conformational changes and that the S2’ proteolysis does not occur in the S prefusion state (Fig. 7). This hypothesis is favored by the packing of the S2’ cleavage site next to S2 HR1. If HR1 were to extend towards the pre-hairpin intermediate, this may remove restrictions on the conformation of the fusion peptide and adjacent S2’ cleavage site to allow for this secondary cleavage. Possible additional factors may include the presence of apposing membranes, the shedding of the S1 subunits or the involvement of yet to be identified host factors.

**Figure 7:**
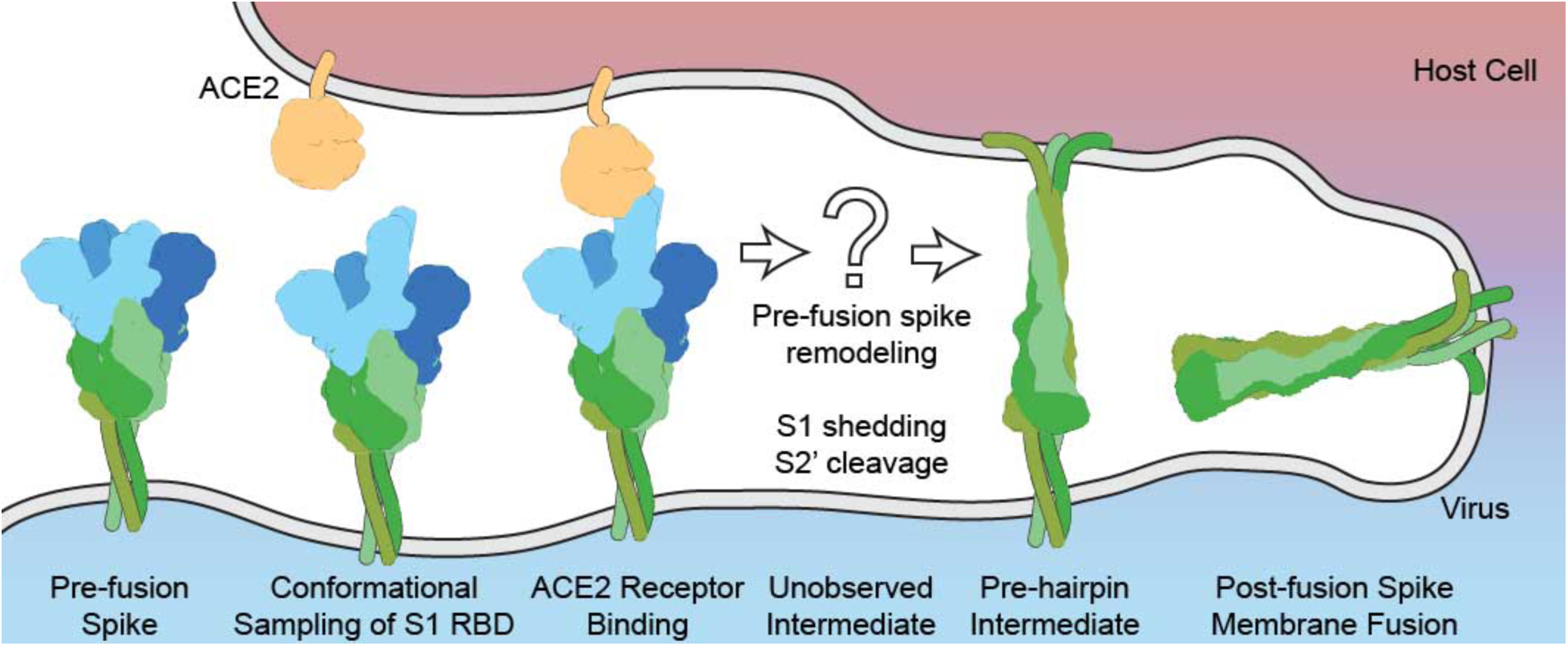
The coronavirus fusion process. The prefusion SARS-CoV S RBD conformationally samples both the down and up states prior to binding the ACE2 receptor. We hypothesize that there is an unobserved intermediate which is not directly trigged by either S1/S2 protease cleavage or ACE2 binding that exposes the S2’ cleavage site. Cleavage at the S2’ site allows membrane insertion of the fusion peptide and subsequent adoption of the postfusion S conformation and membrane fusion. Colors are as in Fig. 3.

## Methods

### Plasmid construction, Protein expression and purification

A mammalian-codon-optimized gene encoding SARS-CoV S (Tor2 strain) residues 1– 1190 with a C-terminal T4 fibritin trimerization domain, an HRV3C cleavage site, an 8xHis-tag and a Twin-Strep-tag was synthesized and subcloned into the eukaryotic-expression vector pαH. Proline-substituted variant harboring K968P and V969P mutations was generated based on this construct. The resulting plasmids, designated SARS S WT and SARS S 2P respectively, were transfected into 1 L FreeStyle 293-F cells (Life Technologies). 293-F cells were used without validation and were not tested for mycoplasma. Three hours after transfection, kifunensine was added to a final concentration of 5 µM. Cultures were harvested after 6 d, and protein was purified from the supernatant using Strep-Tactin resin (IBA). HRV3C protease (1% wt/wt) was added to the protein and the reaction was incubated overnight at 4 °C. The digested protein was further purified using a Superose 6 16/70 column (GE Healthcare Life Sciences).

A gene encoding human ACE2 residues 1-615 with an HRV3C cleavage site, an 8xHis-tag and a Twin-Strep-tag was synthesized and subcloned into the eukaryotic-expression vector pαH. This protein was expressed as described above for S proteins. The HRV3C digested ACE2 protein was further purified using a Superdex 200 column (GE Healthcare Life Sciences).

To make complexes of SARS S 2P with soluble ACE2, the two proteins were combined in a 1:1.1 molar ratio prior to overnight incubation and size-exclusion chromatography on a Superose 6 Increase 10/300 (GE Healthcare Life Sciences).

### Electron microscopy data collection

3 µL of 0.47 mg/mL of protein or protein complexes were mixed with 1 µL of 0.04 % (w/v) amphipol A8-35 (Anatrace) just prior to grid preparation. 3 µL of the protein mixture, final concentration 0.35 mg/mL, was spotted on to plasma cleaned C-flat grids with a 2 µm/2 µm spacing. Grids were blotted for 3 seconds before plunge freezing in liquid ethane on a Vitrobot. Grids were loaded onto a Titan Krios and data was collected using Leginon^32^ at a total dose of 65 e^-^/Å^2^. Frames were aligned with MotionCor2 (UCSF)^33^ implemented in the Appion workflow^34^. Particles were selected using DoG picker^35^. Images were assessed and particle picks were masked using EM Hole Punch^36^. The CTF for each image was estimated using Gctf^37^.

### Electron microscopy data processing

Initial particle stacks were cleaned using multiple rounds of 2D classification in RELION^38^. Good particles were selected as resembling prefusion coronavirus spikes. For the SARS S 2P and trypsin-treated SARS S 2P, all particles from the clean stacks were used for reconstruction with C3 symmetry. All datasets were extensively sorted using 3D classification to examine heterogeneity in the S1 RBDs as described previously^17^. Briefly, 3D masks were defined to encompass the possible heterogeneity at each S1 RBD position. The density within these masks was then removed from unfiltered, unsharpened reconstructions. We then used relion_project with image subtraction to create a particle stack containing only the signal arising from the masked density. Finally, we used focused 3D classification to identify compositional and conformational states at each S1 RBD position. All 3D reconstructions were produced with RELION^38^ and final refinements were performed with a six-pixel soft-edge solvent mask. Post-processing was applied to each reconstruction to apply B-factor sharpening and amplitude corrections as well as to calculate local resolution maps.

Coordinate models were built for several of the high-resolution reconstructions using 5I08.pdb^16^, 2AJF.pdb^23^ and 5X4S.pdb^22^ as template models with reference to a recently published wild-type SARS S ectodomain (5X58.pdb^22^). Manual model building was preformed in Coot^39^ with real-space refinement refinement in Rosetta^40^ and PHENIX^41^ with final rounds of refinement performed in PHENIX.

### Limited proteolysis

10 µg of either SARS-CoV S 2P or SARS-CoV S WT ectodomains were mixed with soluble ACE2 in a 1:1.1 ratio and incubated overnight at 4˚C. Proteins were digested with 0.1% (w/w) TPCK trypsin at room temperature. For SDS-PAGE analysis, samples were removed at indicated time points, immediately mixed with SDS loading buffer and incubated at 95˚C for 1 minute. For EM analysis, the proteolysis reaction was stopped after two hours using phenyl-methyl-sulfonyl-floride (PMSF) at a final concentration of 1 mM and then immediately loaded onto a Superose 6 Increase 10/300 size-exclusion column.

### Data availability

Plasmid DNA used in this study is available upon request. Electron microscopy maps are deposited in the Electron Microscopy Data Bank with accession codes: 7573, 7574, 7575, 7576, 7577, 7578, 7579, 7580, 7581, 7582, 7584, 7585, 7586, 7601, 7602, 7603, 7604, 7605, 7606, 7607 and 7608. Atomic models for select maps are deposited in the Protein Data Bank with accession codes 6CRV, 6CRW, 6CRX, 6CRZ, 6CS0, 6CS1 and 6CS2.

## Acknowledgements

We gratefully acknowledge Travis Nieusma, Charles Bowman, Jean-Christophe Ducom and Bill Anderson for microscopy and computational support. We also thank Lauren Holden for a critical reading of this manuscript. This work was supported by grants from NIH/NIAID to A.B.W and J.S.M. (AI127521) and to R.N.K (AI123498). Computational resources for electron microscopy at The Scripps Research Institute are supported by NIH grant OD021634.

## Author Contributions

R.N.K, J.S.M and A.B.W conceived and designed experiments. N.W. and D.W. expressed and purified protein. R.N.K. and H.L.T. collected electron microscopy data. R.N.K. and J.P. processed electron microscopy data. R.N.K and C.A.C. built and refined atomic models. B.S.G. and K.S.C. contributed support and advice. All authors contributed to writing the manuscript.

## Competing financial interests

R.N.K, N.W., J.P., H.L.T., C.A.C, K.S.C, B.S.G., J.S.M., and A.B.W. are inventors on US patent application no. 62/412,703, entitled “Prefusion Coronavirus Spike Proteins and Their Use.”

